# Primordial capsid and spooled ssDNA genome structures penetrate ancestral events of eukaryotic viruses

**DOI:** 10.1101/2021.03.14.435335

**Authors:** Anna Munke, Kei Kimura, Yuji Tomaru, Han Wang, Kazuhiro Yoshida, Seiya Mito, Yuki Hongo, Kenta Okamoto

**Affiliations:** The Laboratory of Molecular Biophysics, Department of Cell and Molecular Biology, Uppsala University, Uppsala, Sweden; Department of Biological Resource Science, Faculty of Agriculture, Saga University, Saga, Japan; Fisheries Technology Institute, Japan Fisheries Research and Education Agency, Hatsukaichi, Hiroshima, Japan; Graduate School of Agriculture, Saga University, Saga, Japan; Bioinformatics and Biosciences Division, Fisheries Resources Institute, Japan Fisheries Research and Education Agency, Fukuura, Kanazawa, Yokohama, Kanagawa, Japan

## Abstract

Marine algae viruses are important for controlling microorganism communities in the marine ecosystem and played fundamental roles during the early events of viral evolution. Here, we have focused on one major group of marine algae viruses, the ssDNA viruses from the *Bacilladnaviridae* family. We present the capsid structure of the bacilladnavirus, *Chaetoceros tenuissimus* DNA virus type II (CtenDNAV-II), determined at 2.3 Å resolution. A structure-based phylogenetic analysis supported the previous theory that bacilladnaviruses have acquired their capsid protein via horizontal gene transfer from a ssRNA virus. The capsid protein contains the widespread virus jelly-roll fold, but has additional unique features; a third β-sheet and a long C-terminal tail. Further, low-resolution reconstructions of the CtenDNAV-II genome revealed a partially spooled structure, an arrangement previously only described for dsRNA and dsDNA viruses. Together, these results exemplify the importance of genetic recombination for the emergence and evolution of ssDNA viruses and provide important insights into the underlying mechanisms that dictate genome organisation.

## Introduction

Marine algae viruses prevail massively in the oceans and greatly affect the global ecosystem by causing mortality and lysis of microbial communities, releasing organic carbon and other nutrients back into the environment (the ‘viral shunt’), thereby affecting the dynamics of algal blooms, the global oxygen level, and the marine nutrient and energy cycling (Fuhrman, 1999; Suttle, 2007; Wilhelm and Suttle, 1999). The viruses typically have a very narrow host range, thus causing host-specific mortality and control of algae host populations (Brussaard, 2004).

Virus capsid evolution is an increasingly growing research field that rely both on sequence and structural analysis, as well as on computational approaches (Abrescia et al., 2012; Kazlauskas et al., 2017; Krupovic and Koonin, 2017; Munke et al., 2020; Nasir and Caetano-Anollés, 2017; Okamoto et al., 2020, 2016; Wang et al., 2015). These studies range from narrow comparisons between specific virus groups (Kazlauskas et al., 2017; Munke et al., 2020; Okamoto et al., 2020, 2016) to large-scale examinations across the entire virosphere (Krupovic and Koonin, 2017) and have contributed to the identification of unique structural traits among certain viruses (Munke et al., 2020; Okamoto et al., 2020, 2016; Wang et al., 2015) and the revelation of evolutionary relationships between seemingly unrelated viruses (Abrescia et al., 2012; Diemer and Stedman, 2012a; Kazlauskas et al., 2017; Nasir and Caetano-Anollés, 2017). Moreover, a defined number of viral lineages have been identified based on capsid protein folds (Abrescia et al., 2012; Nasir and Caetano-Anollés, 2017), which were originally acquired from cells on multiple independent occasions (Krupovic and Koonin, 2017). Since unicellular marine organisms were the earliest eukaryotes on earth, they were presumably host of the most ancient viruses (Dolja and Koonin, 2018; Koonin et al., 2015). Present-day viruses infecting unicellular organisms, such as unicellular algae, therefore likely retain genetic and structural characteristics from their ascendant (Koonin and Dolja, 2014; Munke et al., 2020), and are consequently an essential group of viruses for interrogating viral capsid evolution.

Bacilladnaviruses is one of the major group of viruses infecting eukaryotic algae. They carry a circular ssDNA genome of ∼6 kb, which is partially double-stranded (∼700-800 bp) and encodes three proteins; one coat protein, one replication protein (Rep), and a third protein with unknown function (Kimura and Tomaru, 2015a). Until recently, bacilladnaviruses have been included in the informal group CRESS DNA viruses (for *circular Rep-encoding ssDNA viruses*), but this group has now formed the phylum *Cressdnaviricota* based on their relatively conserved *Rep* protein (Krupovic et al., 2020). Contrary, the capsid proteins of CRESS DNA viruses are very diverse and have presumably been acquired from RNA viruses on multiple independent occasions (Diemer and Stedman, 2012a; Kazlauskas et al., 2017; Krupovic, 2013; Roux et al., 2013; Tisza et al., 2020). Specifically, bacilladnaviruses were suggested to have acquired their capsid proteins through horizontal gene transfer (HGT) from an ancestral noda-like virus (Kazlauskas et al., 2017). This is not an unreasonable scenario considering the prevalence of noda-like viruses in the aquatic environment (Wolf et al., 2020). However, structural evidence has until now been pending.

Here, we present 3D reconstructions that reveal both the capsid and genome organization of the bacilladnavirus *Chaetoceros tenuissimus* DNA virus type II (CtenDNAV-II) (Kimura and Tomaru, 2015a). An atomic model of the capsid protein could be constructed from the 2.3 Å resolution capsid structure. Structure-based phylogeny was used to demonstrate that the capsid of bacilladnaviruses indeed is structurally more similar to capsids of RNA viruses than to those of other ssDNA viruses, corroborating the HGT theory in early virus evolution. In addition, low-resolution density maps of the ssDNA genome suggest a partially spooled genome packaging mechanism, which has previously only been described for dsRNA and dsDNA viruses.

## Results

### Summary of structure determination

The structure of the CtenDNAV-II viron was determined using cryo-electron microscopy (cryo-EM). The capsid was reconstructed by imposing icosahedral symmetry (I4) and using 33,507 particles to an overall resolution of 2.3 Å using the “gold standard” Fourier shell correlation (FSC) 0.143 criterion (Henderson et al., 2012; Scheres and Chen, 2012) (Supplementary Fig. 1C). The local resolution of the capsid reconstruction was distributed between 2.3 and 9.1 Å and estimated using ResMap (Kucukelbir et al., 2014) (Supplementary Fig. 1D). An atomic model of the capsid was built, refined, and validated according to the cryo-EM map. During 3D classification it became apparent that possibly also the capsid interior possessed higher order structure, yet no density was observed in the final icosahedrally averaged high-resolution reconstruction. However, by employing a previously described method of subtracting the contribution of the capsid (Ilca et al., 2019), two low-resolution genome reconstructions were obtained by using a subset of 21,559 particles and without imposing any symmetry (C1). The resolution of these two reconstructions were automatically estimated by Relion according to the FSC = 0.143 criterion to 13 and 23 Å. Supplementary Fig. 2 shows the FSC curves of the C1 reconstructions, where also the resolution criterion FSC = 0.5 (20 and 26 Å) is indicated as a comparison. Data acquisition and processing, refinement, and validation statistics are summarized in Supplementary Tables 1 and 2.

### Different conformations and unmodelled density blob within the asymmetric unit

The CtenDNAV-II capsid displays T = 3 symmetry, i.e. 180 capsid protein protomers assemble such that the asymmetric unit comprises 3 capsid subunits in 3 quasi-equivalent positions termed A, B, and C (Figs. 1A and B). For subunit A residues 64-72 and 77-371 were modelled, for subunit B residues 64-371 were modelled, and for subunit C residues 60-365 and 378-384 were modelled (Fig. 2C). Apart from residues 73-76 that were unmodelled in subunit A, the A- and B-subunits were close to identical (Fig. 1C): the C-terminus of the A- and B-subunits form long tails that end on the capsid surface around the 3- and 5-fold axes respectively. Here, the last 19 residues could not be modelled (Fig. 2C), however, additional density was visible in the map when the contour level was decreased (Supplementary Fig. 3A), indicating that the C-terminus forms flexible protrusions on the capsid surface. The termini of subunit C differs from the A- and B-subunits (Fig. 1C). Contrary to the long tail towards the capsid surface, the C-terminus of the C-subunit is directed towards the capsid interior (Fig. 1C). The N-termini of all subunits are presumably located on the capsid interior, however, the first 59-63 residues could not be modelled (Fig. 2C). Numerous positively charged residues can be found in the N-terminus, thus possibly interacting with the genome. Residues 366-385 were only partially modelled and the last residues (386-390) were unmodelled in all subunits (Fig. 2C), suggesting different conformations and/or flexibility. The N-terminus of the C-subunit has a small α-helix on the capsid interior that was not present in the other two subunits (Fig. 1C). The modelling of the C-subunit was aided by a structure prediction (Supplementary Fig. 4) from the AlphaFold package (Jumper et al., 2021). The termini of the predicted model were more similar to the C-subunit than to the A- and B-subunits. The N-terminus of the AlphaFold model was confidently predicted (pLDDT >90) from H60, which was also the first residue of the C-subunit that could be confidently modelled based on the cryo-EM map (Fig. 2C). As expected, based on the experimental data, the first 59 residues of the N-terminus of the predicted structure are disordered. The C-subunit was confidently modelled based on the cryo-EM map until R365 (Fig. 2C). With guidance from the AlphaFold model, another seven residues (D378-F384) could be modelled. Residues 366 to 377 could not be confidently modelled based on the experimental data, and are thus not included in the final model (PDB 7NS0), however based on the appearance of the cryo-EM density and the predicted model that has a pLDDT score ranging between 77 and 93 in this part, an α-helix is likely located there (Supplementary Fig. 4). The modelled termini and corresponding cryo-EM map are displayed in Fig. 1D.

**Figure 1.**
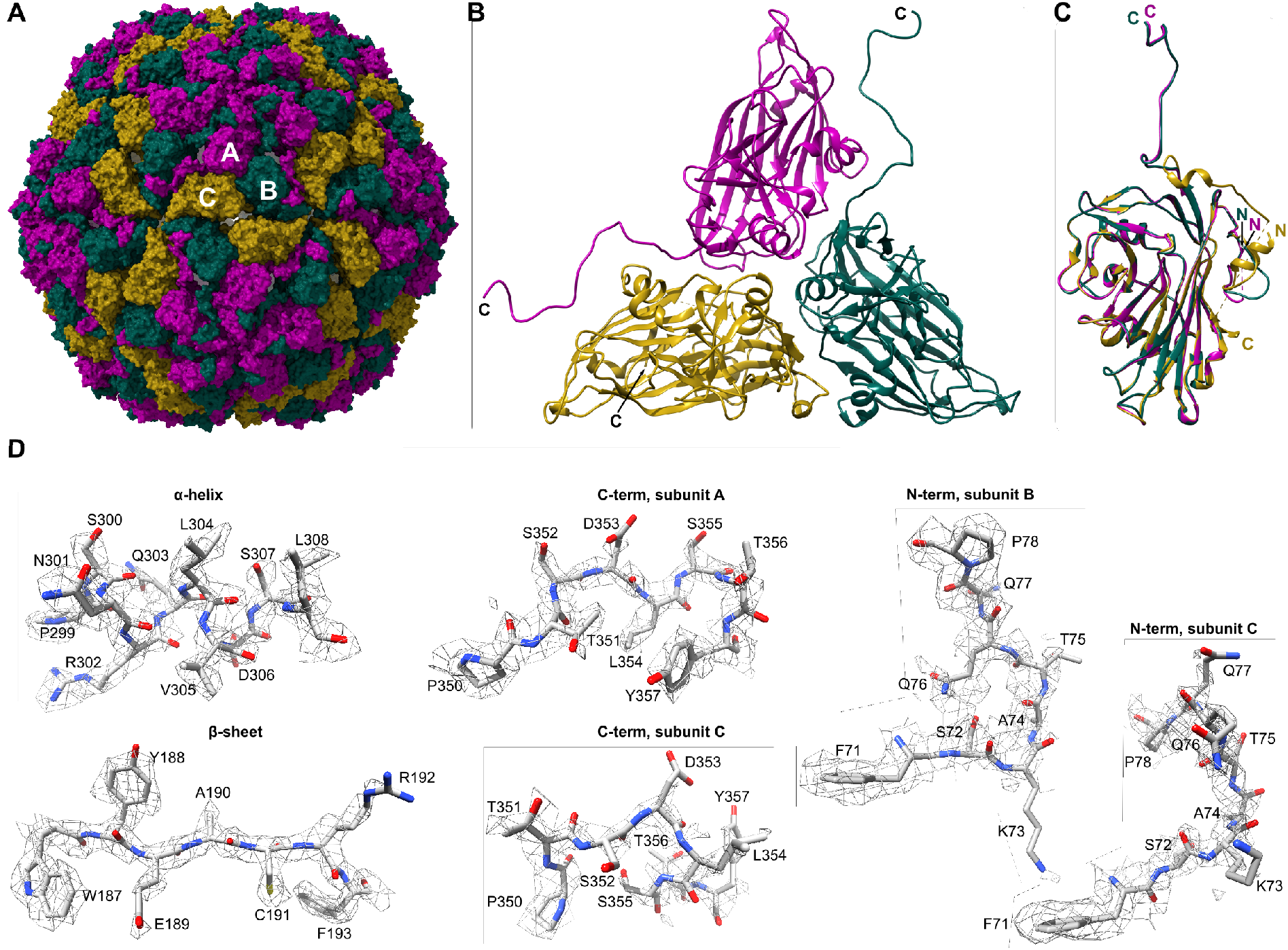
Atomic model of the CtenDNAV-II capsid. The three subunits, A, B and C, are coloured purple, green and yellow, respectively, in panel A-C. (**A**) The entire capsid rendered with a surface representation viewed down an icosahedral two-fold axis. (**B**) The secondary structure of one single asymmetric unit viewed from the outside. (**C**) Superimposition of the three subunits. (**D**) Refined side chains of representatives of secondary structural elements and areas that differ between the three subunits.

**Figure 2.**
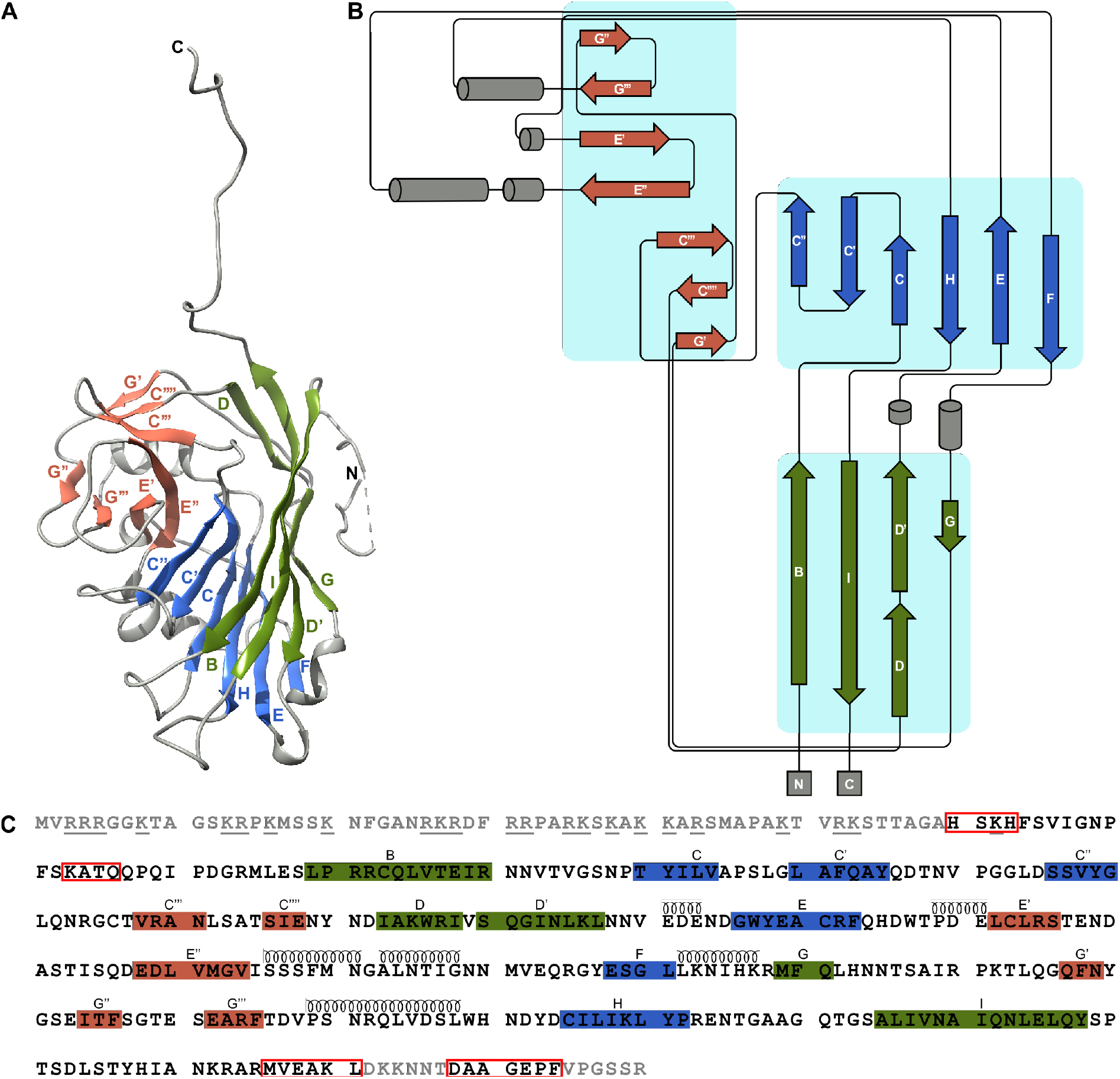
Capsid protein topology and structural organisation. The β-strands are coloured according to which β-sheet they belong to. The β-strands within the green and blue β-sheets are named alphabetically according to the conventional jelly-roll fold nomenclature (B to I) (**A**) The secondary structure of subunit A. (**B**) Schematic diagram of the secondary structure. (**C**) The amino acid sequence of the CtenDNAV-II capsid protein, starting from residue 1. Each line has 70 residues and is further subdivided into blocks of 10 residues by spaces within the sequence. The residue numbering is the same as in PDB entry 7NS0. Modelled and unmodelled residues are coloured black and grey, respectively. Residues highlighted with red rectangles were partly unmodelled: H60-H63 in subunits A and B, F73-P76 in subunit A, M366-L371 in subunit C, and D378-F384 in subunits A and B. The assigned secondary structure is shown schematically above the sequence. The underlined residues in the unmodelled N-terminus indicate the numerous positively charged residues

An unmodelled density blob was visible in the interface of the three subunits that constitute one protomer (Supplementary Fig. 5A). Metal ions, commonly Ca^2+^ (Fisher and Johnson, 1993; Hogle et al., 1983) but also Zn^2+^ (Mathieu, 2001) are sometimes found in the interface of capsid protein subunits and have shown to be important for viron stability. The unmodelled density in the subunit interface of CtenDNAV-II has a pyramidal to spherical shape (Supplementary Fig. 5C) and is surrounded by six arginine residues (R84 and R270) that resemble an octahedral coordination geometry (Supplementary Fig. 5A). The distance from the centre of the density to surrounding nitrogen atoms on the arginine sidechains is 3.7-4.0 Å. The long distance and the fact that arginine residues would be in the coordination sphere suggest that a metal ion at this position is unlikely (Dokmanić et al., 2008). However, a calcium ion has been reported in nodaviruses (Johnson et al., 2001) at a close to identical location (2.6 Å apart) to the unmodelled blob in CtenDNAV-II (Supplementary Fig. 5B), which is interesting from an evolutionary perspective as described later. Considering the location of the unmodelled density, close to the inside of the capsid (Supplementary Fig. 5C), another possibility could be that it originates from a piece of genome. The first residue (F64) to be modelled in all subunits of the positively charged N-terminus is also located close to the subunit interface (yellow stars in Supplementary Fig. 5A), further indicating that this area is important for genome interaction. However, the unmodelled blob appears isolated and structurally obstructed by R84 from the capsid interior (Supplementary Fig. 5C). A multiple sequence alignment (Supplementary Fig. 5D) shows that a positively charged residue (either arginine or lysine) is rather common (16 out of 24 sequences) among bacilladnaviruses at the corresponding position of R84. The second arginine, R270, is very unusual at the corresponding position, however several sequences have a histidine one amino acid upstream, which potentially could serve the same function. Future research will have to discern the true nature of the blob.

### Unique features of the CtenDNAV-II capsid protein jelly-roll fold

The canonical viral jelly-roll consists of eight anti-parallel β-strands that are named from B to I and arranged in two four-stranded sheets (BIDG and CHEF). The loops connecting each strand are named BC, CD, etc. (Harrison et al., 1978; Rossmann et al., 1985). For CtenDNAV-II, the two sheets are formed by strands BIDD’G and C’’C’CHEF, respectively (Fig. 2A-C), thus containing three additional strands (D’, C’ and C’’) compared to the standard viral jelly-roll fold. In addition, a third antiparallel β-sheet with seven strands is intertwined within the jelly-roll, i.e. strands from the third sheet are formed by extensions of loops CD, EF and GH of the jelly-roll. The third sheet, which is located on the capsid surface, is thus composed of two C-strands (C’’’ and C’’’’), two E-strands (E’ and E’’), and three G-strands (G’, G’’ and G’’’) (Fig. 2B). Notably, AlphaFold accurately and confidently predicted the jelly-roll fold as well as the fold of the additional β-sheet, which had pLDDT scores of >90 and >70, respectively (Supplementary Fig. 4). In conclusion, the capsid protein of CtenDNAV-II has three unique features: three additional strands in the jelly-roll fold, an extra surface-exposed β-sheet, and an unusual conformation of the C-termini of subunit A and B that extends out from the jelly-roll core as long tails toward the capsid surface.

### Capsid proteins from CtenDNAV-II and ssRNA viruses are similar

Previous structures of so-called CRESS DNA viruses (phylum *Cressdnaviricota*) (Krupovic et al., 2020) include viruses from families *Ciroviridae* (eg. 3R0R and 5ZJU), *Geminiviridae* (eg. 6F2S and 6EK5), and *Nanoviridae* (6S44). The capsid proteins of these three families also contain a jelly-roll domain, but lack the third surface-exposed β-sheet and N-terminal tail found in CtenDNAV-II. The DALI web server (Holm, 2020a) was used as an initial step to investigate the previous theory that ssDNA viruses have acquired their capsid protein from ssRNA viruses (Kazlauskas et al., 2017; Krupovic, 2013). Indeed, the result indicated that the CtenDNAV-II capsid protein is more similar to capsid proteins of ssRNA viruses than to other ssDNA viruses (Supplementary Fig. 6). The closest ssDNA virus was that of Beak and feather disease virus (*Circoviridae*), which ended up on 11th place (z-score 8.4) behind ten RNA viruses. Highest similarity was found between CtenDNAV-II and ssRNA viruses from families *Carmotetraviridae, Alphatetraviridae* and *Nodaviridae*, which had z-scores of 14.4-15.1 (Supplementary Fig. 6). Notably, only nodaviruses from the genus *Alphanodavirus* were included and none from *Betanodavirus* or the informal group of gamma-nodaviruses. Z-scores below 2 are considered insignificant, scores between 2 and 8 are a grey zone, scores between 8 and 20 indicates that two structures probably are homologous, and with a z-score above 20 they are definitely homologous. For an in-depth explanation of the DALI method and z-scores, see Holm (2020) (Holm, 2020b).

A structure-based phylogenetic analysis was used to further investigate the relationship between the capsid proteins of CtenDNAV-II and ssRNA viruses. In summary, structures with DALI-scores >8 and three additional CRESS DNA viruses were used together with the CtenDNAV-II capsid protein structure as inputs to the program MUSTANG (Konagurthu et al., 2006). A complete list of structures used for the analysis is shown in Figure 3–figure supplement 1. The Root Mean Square Deviation (RMSD) values provided by MUSTANG were then used to create phylogenetic trees (Fig. 3 and Figure 3–figure supplement 2). The RMSD matrices created by MUSTANG that were used to create the phylogenetic trees are available in Figure 3–Source Data 1 and 2. The compared structures have various polypeptide lengths, ranging from 172 amino acids for Faba bean necrotic stunt virus (*Nanoviridae*) to 644 amino acids for Nudaurelia capensis omega virus (*Alphatetraviridae*), and thus the longer structures e.g. from CtenDNAV-II, carmotetraviruses, alphatetraviruses, nodaviruses and birnaviruses have additional folds and domains in addition to their jelly-roll fold core (Supplementary Fig. 6). The phylogenetic analysis was therefore performed using only the jelly-roll fold core, for which the result is presented in Figure 3. The selected amino acid numbers for each structure in the jelly-roll fold comparison are listed in Figure 3–figure supplement 1. Figure 3 displays three clades with different ssDNA viruses in each clade. CtenDNAV-II is placed together with RNA viruses from families *Nodaviridae, Alphatetraviridae, Carmotetraviridae* and *Birnaviridae* (Clade 1), thus corroborating the result from the DALI search (Supplementary Fig. 6). The ssDNA circoviruses were organised in a second clade together with ssRNA viruses from families *Tombusviridae, Hepaviridae* and *Solemoviridae*, whereas the other two ssDNA families *Geminiviridae* and *Nanoviridae* were placed alone in the third clade. As a comparison, the same analysis was performed by using full-length chains (Figure 3–figure supplement 2). The result is to some extent similar: CtenDNAV-II and nodaviruses still share structural similarities, and CtenDNAV-II and other ssDNA viruses are organised into different clades, however the details in the tree differ between the two approaches. For example, only CtenDNAV-II and the nodaviruses appear related, whereas carmotetraviruses, alphatetraviruses, and birnaviruses are placed in the other clades. However, our opinion is that Figure 3 better represents the evolutionary relationship between these viruses, since the jelly-roll fold, which is the conserved structural core, in some cases are misaligned when using the full-length chains (Figure 3–figure supplement 3-12).

**Figure 3.**
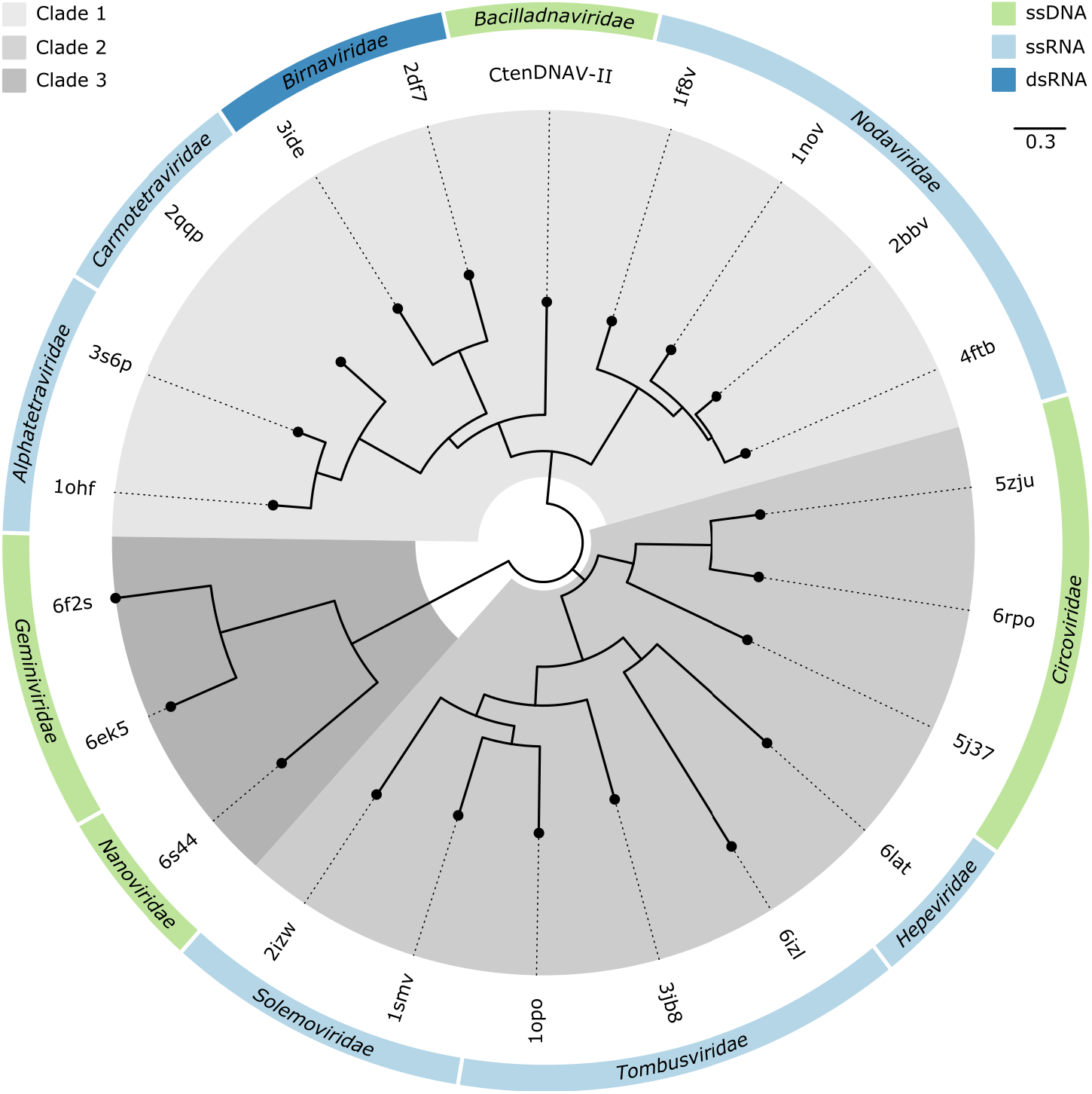
Structure-based phylogenetic tree of the jelly-roll folds. The three major clades are highlighted in different shades of grey. The tree is reconstructed based on RMSD values from superpositions performed on the common jelly-roll fold core. The chain identity and residues included for each structure are listed in Figure 3–figure supplement 1. A phylogenetic tree on full-length chains is shown in Figure 3–figure supplement 2. The superpositions between CtenDNAV-II and the other structures are displayed in Figure 3–figure supplement 3-12, with one representative structure for each virus family. The RMSD tables of the jelly-roll fold and full-length superpositions are available in Figure 3–Source Data 1 and 2, respectively.

In conclusion, our structural analysis revealed that the CtenDNAV-II capsid protein is more similar to capsid proteins of RNA viruses than to other ssDNA viruses (Fig. 3), corroborating the previous HGT theory by Kazlauskas et al (Kazlauskas et al., 2017). Further, the ssDNA viruses were organised into three different clades, thus supporting a polyphyletic origin of ssDNA viruses and the independent acquisition of capsid protein genes from different RNA viruses (Krupovic, 2013).

The jelly-roll fold of CtenDNAV-II align well with the other jelly-roll folds in Clade 1 (Figure 3– figure supplement 3-6), with RMSD values of 1.4-1.7 (Figure 3–Source Data 1), compared to the jelly-roll folds in the other clades (Figure 3–figure supplement 7-12) that had RMSD values of 2.2-3.6. As expected, the differences between the capsid proteins from CtenDNAV-II and carmotetraviruses, alphatetraviruses, nodaviruses, and birnaviruses lie in areas that are either surface exposed or located on the capsid interior (Fig. 4), i.e. structural elements that have evolved based on the host and on genome packing requirements, respectively. All capsid proteins in Clade 1 have specific surface structures in addition to the jelly-roll fold (Fig. 4 and Fig. 4–figure supplement 1). While the surface structures of CtenDNAV-II and nodaviruses are formed by extensions of loops in the jelly-roll, carmotetraviruses, alphatetraviruses and birnaviruses instead have larger separate domains (called Ig-like and P-domains) inserted between two jelly-roll strands. Unique for CtenDNAV-II is the long C-terminal tail in subunits A and B that also extends to the capsid surface, while the C-termini in the capsid protein of the other viruses can be found on the capsid interior similarly as the C-terminus of the C-subunit in CtenDNAV-II (Fig. 4 and Fig. 4–figure supplement 1). Carmotetraviruses, alphatetraviruses, nodaviruses, and birnaviruses have α-helices located on the capsid interior that are formed by the capsid protein termini, which in some structures have been shown to interact with the viral RNA (Speir et al., 2010). Of the three CtenDNAV-II subunits, the C-subunit shows highest similarity to the capsid proteins of the other viruses in Clade 1: both termini are located on the capsid interior and the termini forms α-helices. The model of the C-subunit has one α-helix in the N-terminus, however, based on the cryo-EM map and the predicted model by AlphaFold, also the C-terminus is likely to form an α-helix (Supplementary Fig. 4). Superpositions of the capsid proteins from families *Carmotetraviridae, Alphatetraviridae, Nodaviridae*, and *Birnaviridae* with the predicted model of the CtenDNAV-II capsid protein shows that the two predicted terminal α-helices align well with α-helices from the experimentally determined structures. Finally, an additional similarity is found in the N-terminus of CtenDNAV-II and the ssRNA viruses in Clade 1. Due to disorder, up to 63 residues of the highly positively charged terminus could not be modelled for CtenDNAV-II (Fig. 2C). Likewise, the first ∼40-70 residues, also heavily populated with arginine and lysine residues, are unmodelled for the ssRNA viruses. This is in contrast to the dsRNA birnaviruses, whose N-terminus is largely modelled (missing 5-10 residues) and less charged.

**Figure 4.**
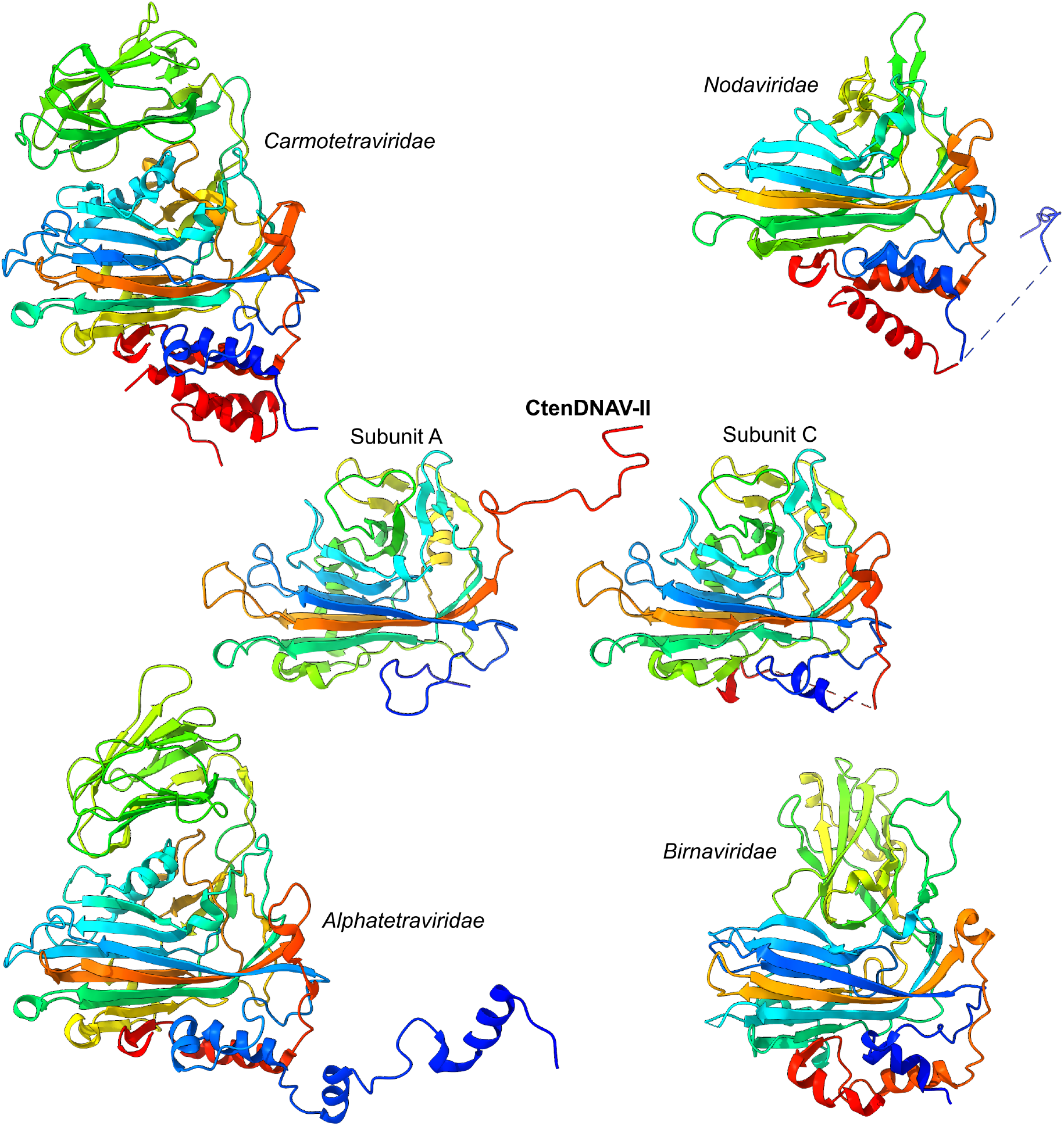
Comparison of capsid proteins in Clade 1. Representatives of capsid proteins in Clade 1 viewed from the side with surface and interior structures on top and bottom, respectively. The same representatives were used as in Figure 3–figure supplement 3-6. The chains are coloured as rainbow from blue (N-terminus) to red (C-terminus). Figure 4–figure supplement 1 shows same images, but coloured according to domain.

### CtenDNAV-II genome is partially spooled

The genome organization of CtenDNAV-II is shown in Figure 5. The CtenDNAV-II genome consists of an ordered outer layer (Fig. 5B and E) that is partially spooled around a disordered core (Fig. 5C). The outer layer (EMDB-12555) was reconstructed at 13-20 Å by masking out the capsid, while the core (EMDB-12556) was reconstructed at 23-26 Å by using a spherical mask (Supplementary Fig. 2) (see Materials and Methods for details). The reconstruction of the outer genome layer displays a coil of three turns that are positioned between the icosahedral 5-fold axes (Fig. 5E). On each side of the three turns are additional DNA fragments that do not follow the same spooling arrangement (Fig. 5B). The spooled genome packaging has previously only been described in viruses with double stranded genomes (Ilca et al., 2019; Liu et al., 2019; Wang et al., 2019), whereas the genomes of ssRNA viruses forms secondary structures that organise into branched networks (Dai et al., 2017; Gorzelnik et al., 2016; Koning et al., 2016) or a dodecahedral cage (Hesketh et al., 2015; Johnson et al., 2001). Below each icosahedral 5-fold axis of the capsid the reconstruction of the outer genome layer displays protrusions towards the capsid (Fig. 5B and E), which could originate from genome and/or the long N-terminus that was unmodelled for the capsid protein (Fig. 2C). Connections between the outer genome layer and the core seem to be confined to two specific areas on opposite sides of the outer layer (Supplementary Fig. 6). Movie 1 shows the asymmetric reconstruction in relation to the capsid.

**Figure 5.**
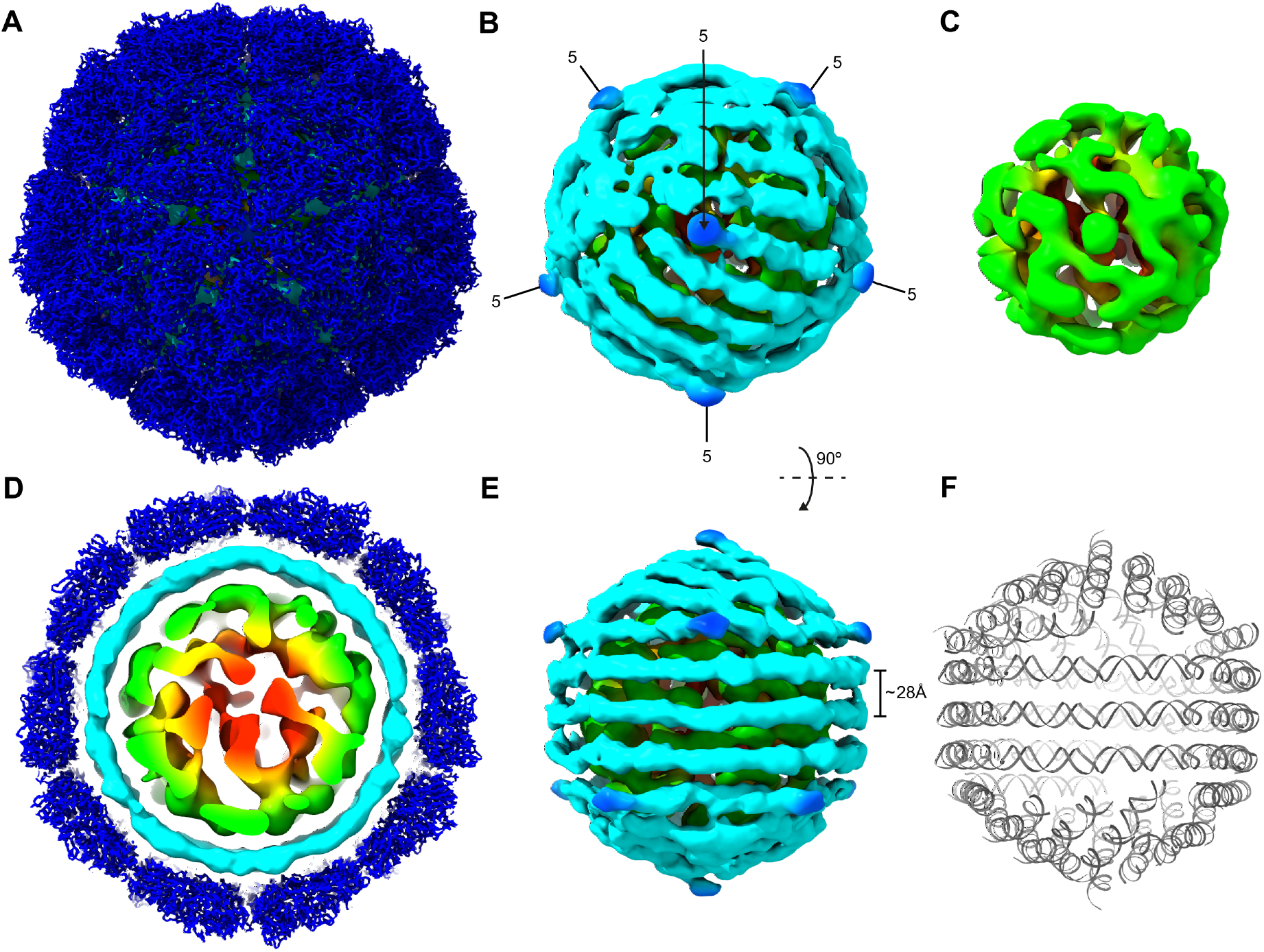
Asymmetric reconstructions of CtenDNAV-II radially coloured from red to blue. (**A**) The capsid (blue) reconstructed with icosahedral symmetry and the asymmetric reconstructions sighted below viewed down the icosahedral five-fold axis. (**B**) Same view as (A) but with the icosahedral reconstruction removed. A protrusion (blue) from the outer genome layer (cyan) is visualised at each five-fold axis (labelled as 5). (**C**) Same view as in (A) and (B) but with the icosahedral and outer layer reconstructions removed. (**D**) A thin slice of the reconstructions shown in (A). (**E**) The picture in (B) has been rotated 90**°**. (**F**) An approximate model of the outer genome layer, same viewing direction as in (E).

Considering that spooled genome arrangements have only been observed among viruses with double-stranded genomes and that the distance between the parallel turns are about 28 Å (Fig. 5E), similar to other double-stranded genomes (Ilca et al., 2019; Wang et al., 2018), the outer genome layer of CtenDNAV-II is likely formed by double-stranded DNA. The low resolution of the outer layer prevented precise modelling; however, an approximate arrangement of the outer layer was achieved by manually combining short (5-20 nt) A-T base pairs of ideal B-form dsDNA into the density (Fig. 5F). CtenDNAV-II carries a covalently closed circular ssDNA genome of 5,770 nucleotides (nt) that includes a partially double-stranded region (669 nt) (Kimura and Tomaru, 2015b). The outer genome layer corresponds to about 50 % of the total CtenDNAV-II genome (see Materials and Methods for details), which further supports that the outer layer is double-stranded, since it seems unlikely that the genome core could accommodate much more than the other 50 %. The partially double-stranded region of the CtenDNAV-II genome is not long enough to occupy the entire outer layer; thus, the outer layer is at least partially composed of ssDNA that forms double-stranded secondary structures.

## Discussion

### Structural insights on eukaryotic jelly-roll fold viruses in evolution

About 30% of all viral capsid proteins adopt the single jelly-roll fold (Krupovic and Koonin, 2017), and the results presented herein provide structural insights on the evolution of this fold. The majority of capsid proteins, including those having the jelly-roll fold, were presumably once acquired from cells on multiple independent occasions (Krupovic and Koonin, 2017). Contrary, the viral replication proteins have no homologous in cellular organisms, and thus, one of the theories to explain the origin of viruses is the so-called ‘chimeric scenario’ where the virus replication genes existed in a pre-cellular world and that the structural proteins were acquired later in evolution from primitive cellular organisms (Krupovic et al., 2019). Further, it has been postulated that the genetic elements in the precellular world were of RNA type (Gilbert, 1986; Krupovic et al., 2019) and that viruses with different genome types can be placed into an evolutionary continuum, from ssRNA, to dsRNA, to ssDNA, and to dsDNA (Krupovic, 2013). Gene exchange between RNA and DNA viruses have been frequently described in the literature. The capsid proteins of CRESS DNA viruses have for example likely been acquired from different RNA viruses (Diemer and Stedman, 2012a; Kazlauskas et al., 2017; Krupovic et al., 2009; Roux et al., 2013). Specifically, the capsid protein gene of bacilladnaviruses were suggested to originate from a primordial noda-like virus (Kazlauskas et al., 2017).

Indeed, our structural analysis suggest that the capsid protein of CtenDNAV-II is structurally more similar to ssRNA and dsRNA viruses (Clade 1) than to other CRESS-DNA viruses (Clade 2 and 3) (Fig. 3), thus corroborating the theory that the jelly-roll capsid proteins of ssDNA, ssRNA and dsRNA viruses originate from the same virus with a primordial jelly-roll fold (Kazlauskas et al., 2017; Krupovic and Koonin, 2017). In addition, the results support the polyphyletic origin of ssDNA viruses and the independent acquisition of capsid protein genes from RNA viruses (Krupovic, 2013). Perhaps the acquisition of the capsid protein gene from a noda-like virus allowed bacilladnaviruses to accommodate a larger genome than other CRESS DNA viruses, such as circoviruses, that forms smaller (<20nm) capsids with T=1 symmetry (Kazlauskas et al., 2017). Our findings also support the previously described close evolutionary relationship between virus families *Carmoteraviridae, Alphatetraviridae, Nodaviridae*, and *Birnaviridae* (Fig. 3). The capsid proteins of the T=4 tetraviruses have been proposed to be derived from the T=3 nodaviruses, and in particular, carmotetraviruses such as *Providence virus* display a higher degree of similarity to nodaviruses than alphatetraviruses such as *Nudaurelia capensis ω virus*, and are thus supposedly closer to the primordial T=4 virus (Speir et al., 2010). The dsRNA birnaviruses (T=13) presumably have a noda/tetravirus-like ancestor (Coulibaly et al., 2005), which exemplifies the evolutionary link between ssRNA and dsRNA viruses. In turn, an ancestral birna-like virus could have been the precursor of the *Reoviridae* outer capsid layer, whereas the inner layer, that has T=1 symmetry, might have been acquired from an ancestral toti-like virus (Coulibaly et al., 2005). To put our new structural insights into a broader context, we have updated the putative emergence of viruses harbouring different genomes types (Fig. 6).

**Figure 6.**
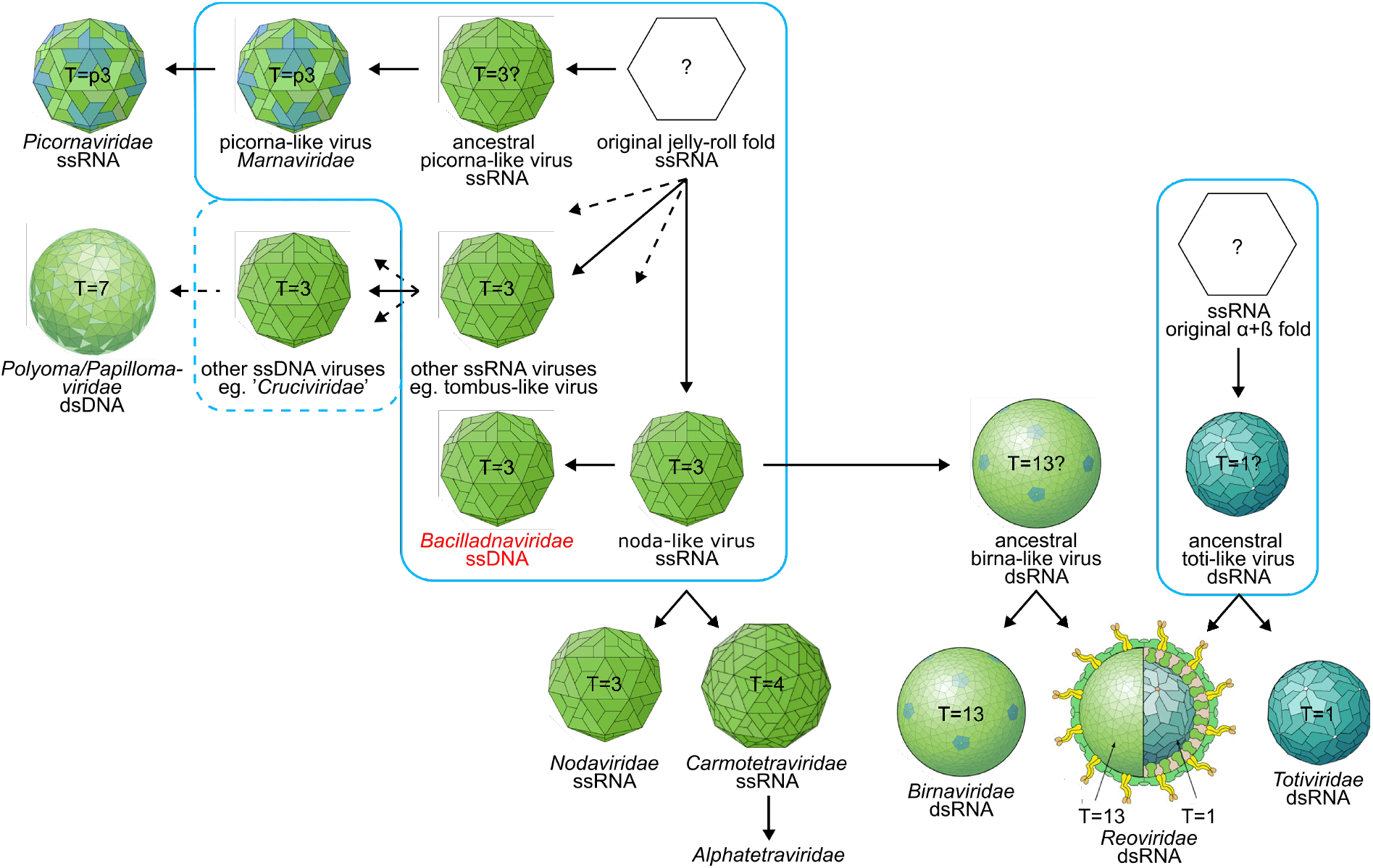
A scheme depicting the putative emergence of virus capsids carrying different nucleic acid genomes. The figure illustrates, based on current knowledge and hypotheses, the most relevant virus groups and the evolutionary relationship between their virus capsids. Events that likely took place at an early stage of evolution in ancient algal pools are circled in blue. The dashed blue line represents that ssRNA and ssDNA virus recombination has occurred at several independent occasions, at different time points in evolution. The figure is based on results described and discussed in this paper as well as previous results and discussions: the relationship between bacilladnaviruses, nodaviruses, carmotetraviruses, alphatetraviruses and binaviruses are described herein as well as in (Coulibaly et al., 2005; Kazlauskas et al., 2017; Krupovic, 2013; Speir et al., 2010); the relationship between the dsRNA viruses are described in (Coulibaly et al., 2005); the evolution of picornaviruses were described in (Munke et al., 2020); the tombus-crucivirus linkage is described in (Diemer and Stedman, 2012a; Roux et al., 2013), and the link between the ssRNA, ssDNA and dsDNA are described by (Kazlauskas et al., 2019; Koonin et al., 2015; Krupovic, 2013). The viron pictures were derived from ViralZone, SIB Swiss Institute of Bioinformatics (Hulo et al., 2011) (https://viralzone.expasy.org/) licensed under a Creative Commons Attribution 4.0 International License.

The structural comparison between CtenDNAV-II and the other viruses in Clade 1 (Fig. 4) revealed the acquired structural features for each virus family. These include for example the Ig-like domain of tetraviruses and the P-domain of birnaviruses, which have been described as putative receptor-binding domains of those viruses (Coulibaly et al., 2005; Garriga et al., 2006; Helgstrand et al., 2004; Penkler et al., 2016; Speir et al., 2010). The unique structural features of CtenDNAV-II included a third β-sheet and a long C-terminal tail of the A and B subunits that extend to and forms flexible protrusions on the capsid surface (Fig. 2A and Supplementary Fig. 3). Similar to the RNA viruses, these acquired features of CtenDNAV-II could potentially be important for host recognition, however future studies will have to confirm this theory. The variable host-specific features also justify why the evolutionary relationship between CtenDNAV-II and the other viruses in this study were deduced based on the MUSTANG analysis using the jelly-roll fold alone (Fig. 3) instead of using the entire chain (Fig. 3– figure supplement 2). The conserved structural element (the jelly-roll fold) is likely to better represent the primordial fold. Contrary to the A and B subunits, the C-terminus of the C-subunit was located on the capsid interior similar to the other viruses in Clade 1 (Fig. 4). In addition, both termini likely forms α-helices (Supplementary Fig. 4) that are structurally well-aligned with those of the RNA viruses. More and longer helices are found among the RNA viruses (Fig. 4) and the two below the C-subunit could thus be remnants from the noda-like virus ancestor, whereas the C-terminus in the A and B subunits has evolved to form other structural elements as a host adaptation. Thus, the C-subunit of CtenDNAV-II presumably resembles the primordial bacilladnavirus fold more that the A and B subunits.

The results presented herein supports the theory of an acquisition of the capsid protein gene in a bacilladnavirus ancestor from a ssRNA noda-like virus via a HGT event, thus supporting the so-called RNA-to-DNA jump scenario for the origin of DNA viruses with jelly-roll capsid proteins (Krupovic, 2012). Other alternative scenarios include the independent emergence (i.e. convergent evolution) and the gradual transition scenarios (Krupovic, 2012). While the latter two cannot be completely ruled out, the first scenario is strongly supported for DNA jelly-roll fold containing viruses, for example due the fact that several different DNA viruses, including bacilladnaviruses, apparently contain capsid proteins that are more similar, on both sequence and structural level, to different RNA viruses, while their rolling-circle replication proteins are polyphyletic (Diemer and Stedman, 2012b; Kazlauskas et al., 2017; Krupovic, 2013; Krupovic et al., 2009; Roux et al., 2013; Tisza et al., 2020). Findings supporting that ssRNA and ssDNA recombination is plausible include the fact that nucleic acid packing can be unspecific (Krupovic, 2013; Routh et al., 2012) and that single-stranded genomes have similar persistence lengths (Cuervo et al., 2013). For more detailed discussions on this topic, see Holmes, 2011; Krupovic, 2012; Moreira and López-García, 2009.

### Genome structures – a consequence of capsid adaptations and packing mechanisms

The technological and computational improvements in cryo-EM structure determination have led to an increasing number of virus genome structures during the last decade. However, determining the genome structure within the symmetric icosahedral capsid still remains challenging. By employing a previously described method of subtracting the contribution of the capsid (Ilca et al., 2019), we were here able to demonstrate the first low resolution density map of a ssDNA genome (Fig. 5 and Movie 1).

Previous cryo-EM studies have revealed different types of genome organization for non-enveloped icosahedral viruses that often reflect the physical properties of the genome and the viron assembly mechanisms. Viruses containing double-stranded genomes (dsDNA and some dsRNA) uses a NTP-driven motor to form spooled genome structures within a preassembled capsid (Ilca et al., 2019; Liu et al., 2019; Wang et al., 2019). The spooled genomes are packaged by flexible interactions between the capsid protein and the genome. The interactions are mediated by small contacts with hydrophobic and/or positively charged amino acid residues of their capsid proteins. Segmented dsRNA reoviruses instead form non-spooled or partially-spooled genomes with pseudo-D3 symmetry that interact with the RNA-dependent RNA polymerase (Ding et al., 2019; Liu and Cheng, 2015; Wang et al., 2018; Zhang et al., 2015). Genomes of ssRNA viruses forms secondary structures that can either form a branched network, such as in *Leviviridae* viruses (Dai et al., 2017; Gorzelnik et al., 2016; Koning et al., 2016), or a dodecahedral cage, such as in *Secoviridae* and *Nodaviridae* viruses (Hesketh et al., 2015; Johnson et al., 2001). Contrary to the large double-stranded genomes that arrange within a preformed capsid, the small ssRNA genomes allow a simultaneous and cooperative genome packing and capsid assembly that is governed by specific interactions between the genome and capsid protein (Hesketh et al., 2015; Koning et al., 2016). Less in known about the genomes of ssDNA viruses, however, similar to viruses with other genome types, the genomes of ssDNA viruses are strongly connected to the packing mechanism (Chapman and Rossmann, 1995; Hesketh et al., 2018; Sarker et al., 2016). Previous structural information on the genomes of ssDNA viruses have been limited to short (<10 nt) nucleotide segments, but have revealed specific interactions between the capsid and DNA of single-stranded type (Chapman and Rossmann, 1995; Hesketh et al., 2018; McKenna et al., 1992). Thus, the density map presented here is the first view of a ssDNA genome organisation.

The genomes of ssDNA viruses have the capability to form biologically functional secondary structures similar to ssRNA viruses (Muhire et al., 2014) and since the capsid gene of bacilladnaviruses has been horizontally transferred from ssRNA viruses, a similar structural arrangement of the genome as in ssRNA viruses is imaginable also for ssDNA viruses. However, the structure of the CtenDNAV-II genome is much more similar to the spooled genomes found in double-stranded viruses (Fig. 5 and Movie 1), thus raising the question whether the CtenDNAV-II capsid pre-assembles before packing the genome. Interestingly, rod-shaped virus-like particles have been found together with CtenDNAV-II particles in infected host cells and were suggested to be precursors of mature virons (Kimura and Tomaru, 2015b). Future studies will be needed to unravel the exact mechanisms behind bacilladnaviron assembly and genome packing.

Another question that arises is if and how the ssRNA virus-like capsid protein of CtenDNAV-II has adapted to pack a dsDNA/dsRNA-like spooled genome? While differences in surface features between the viruses in Clade 1 could reflect variations in host specificity, the differences on the capsid interior could possibly reflect variations in genome organisation (Figure 4–figure supplement 1). Structural information on the genomes of the RNA viruses in Clade 1 is limited to short nucleotide segments and the most complete picture of a possible organisation has been gained from the *Nodaviridae* Pariacoto virus whose genome is organised as a dodecahedral cage that accounts for ∼35% of the total genome (Johnson et al., 2001). The RNA segments have been revealed even when the icosahedral symmetry has not been broken during the structure determination, suggesting that noda-, tetra- and birnaviruses interact more specifically with their genome than CtenDNAV-II does and that their genomes are organised structurally different. No DNA segments were found inside CtenDNAV-II during the capsid reconstruction. Yet, similarities exist in the capsid protein interior domains of CtenDNAV-II and viruses from families *Carmotetraviridae, Alphatetraviridae, Nodaviridae*, and *Birnaviridae* (Figure 4– figure supplement 1). The α-helical domain found in the RNA viruses is not as pronounced in CtenDNAV-II since only the C-subunit has both its termini on the capsid interior, however the presumably two α-helices that exist, based on experimental and predicted models (Supplementary Fig. 4), align structurally well with α-helices of the RNA viruses. In addition, the long disordered and positively charged N-terminus is shared between CtenDNAV-II and the ssRNA viruses in Clade 1.

Our hypothesis is that the CtenDNAV-II acquisition of the T=3 capsid protein gene from a noda/tetra-like virus ancestor allowed packing a larger genome than other CRESS DNA viruses that forms smaller capsids with T=1 symmetry (Kazlauskas et al., 2017). Further, the internal structural features, the positively charged and largely unordered termini, allowed packing a spooled genome that interact non-specifically with the capsid, and whose structure is dictated by the viron assembly mechanism, possibly as a preformed capsid that internalise the ssDNA genome.

## Materials and methods

### Virus production and purification

CtenDNAV-II was produced as previously described (Kimura and Tomaru, 2015a). The crude virus suspension was loaded onto 15 to 50% (w/v) sucrose density gradients and centrifuged at 24,000 × *rpm* (102,170 × *g*) for 18 h at 4°C (Sw40Ti rotor, Beckman Coulter). The fractions of the sucrose gradient were applied to SDS-PAGE. The VP2 capsid protein fractions were pooled and subjected to centrifugation at 28,000 rpm (139,065 × *g*) for 3 h at 4°C (Sw40Ti rotor, Beckman Coulter). The pellet was resuspended in 50 mM Tris (pH 7.4), 100 mM NaCl, and 0.1 mM EDTA.

### Cryo-EM and data collection

An aliquot (3 μl) of purified CtenDNAV-II virons (10 mg ml^-1^) was deposited onto freshly glow-discharged holey carbon-coated copper grids (Quantifoil R 2/2, 300 mesh, copper) followed by 2 s of blotting in 100% relative humidity for plunge-freezing (Vitrobot Mark IV) in liquid ethane. Images were acquired using a Titan Krios microscope (Thermo Fisher Scientific) operated at 300 kV and equipped with a K2 Summit direct electron detector (Gatan) and an energy filter.

### Image processing and 3D reconstruction

The micrographs were corrected for beam-induced drift using MotionCor2 1.2.6 (Zheng et al., 2017), and contrast transfer function (CTF) parameters were estimated using Gctf 1.06 (Zhang, 2016). The RELION 3.1 package (Zivanov et al., 2018) was used for particle picking, 2D and 3D classifications, *de novo* 3D model generation and refinement. A reconstruction of the capsid was generated in I4 symmetry using 33,507 particles, which were obtained by performing 9 consecutive 2D classification steps. The two genome reconstructions were generated in C1 symmetry using 21,559 particles, which were generated by performing 6 consecutive 3D classifications of the 33,507 particles that were obtained from the 2D classification step. Resolutions were estimated using the gold standard Fourier shell correlation (threshold, 0.143 criterion) (Henderson et al., 2012; Scheres and Chen, 2012). The data set and image processing are summarized in Table 1.

To reconstruct the genome without icosahedral symmetry a similar procedure to what has been described by Ilca et al (Ilca et al., 2019) was carried out. The contribution of the capsid was first subtracted from the map created during the final iteration of the I4 refinement job using the Particle subtraction function in Relion. To create a mask for the particle subtraction, the capsid model was first transformed to a density map using the molmap command in Chimera and then an inverted soft edged mask was created from the density map using relion_mask_create with the --invert option. The subtraction was followed by 3D classification (C1 symmetry), which generated the subset of 21,559 particles that was used for the final C1 refinement. The 3D classifications revealed clear density of an outer layer, and an additional subtraction was therefore performed using a spherical mask of 100 Å before the final refinement. The spherical mask was created using relion_mask_create with the --denovo and --outer_radius options. The two maps (the capsid density created by molmap and the circular map) were combined using Chimeras vop command before creating a new inverted mask using relion_mask_create, which was used for subtraction before the final 3D refinement. To reconstruct the core alone the spherical mask was used for the subtraction procedure.

### Model building and refinement

The atomic model of CtenDNAV-II capsid protein was manually built into the density map using Coot (Emsley and Cowtan, 2004). The model was further improved through cycles of real-space refinement in PHENIX (Adams et al., 2010) with geometric and secondary structure restraints, and subsequent manual corrections by Coot were carried out iteratively. Refinement and validation statistics are summarised in Supplementary Table 1. The model of the outer genome layer was constructed by manually combining short (5-20 nt) A-T base pairs of ideal B-form dsDNA into the density using UCSF Chimera X (Goddard et al., 2018). The structure prediction was carried out using AlphaFold 2.0 (Jumper et al., 2021).

### Structure analysis

Structural comparison of the CtenDNAV-II capsid protein was initially carried out by the DALI web server as a heuristic search against all structures (as of 2021-07-05) in the PDB (Holm, 2020a). Unique viruses (i.e. not unique PDB entries) from the DALI search with z-scores >8 were used for structure-based multiple alignments using the program MUSTANG (Konagurthu et al., 2006). In addition, three other CRESS DNA viruses with known structure (6S44, 6EK5 and 6F2S) were included in the MUSTANG analysis. Detailed information on the structures used for the comparison can be found in Figure 3–figure supplement 1. The RMSD values provided by MUSTANG (see Figure 3–Source Data 1 and 2) were used to create phylogenetic trees using the Neighbor-joining method (Saitou, 1987) in MEGA X (Kumar et al., 2018; Stecher et al., 2020), which were further aesthetically modified with FigTree. Figures were prepared using UCSF Chimera X (Goddard et al., 2018).

The number of nucleotides modelled in the coil of three turns (1,260 nt) served as basis for estimating how much of the genome was ordered in the outer layer. UCSF Chimera (Pettersen et al., 2004) was used to calculate the volume of the density corresponding to the three turns, which was compared to the total volume of the outer layer. The outer genome layer was estimated to comprise 3,283 nt, which is 51 % of the CtenDNAV-II genome (5,770 + 669 nt).

### Rich media files

**Movie 1**. Asymmetric reconstruction of the genome in relation to the capsid.

## Acknowledgements

The data were collected at the Cryo-EM Swedish National Facility funded by the Knut and Alice Wallenberg, Erling Persson Family, and Kempe Foundations, SciLifeLab, Stockholm University and Umeå University. We thank Julian Conrad and Dustin Morado for help with data collection. We also want to thank Afonso Vieira for valuable discussions on the CtenDNAV-II capsid structure.

