## Supplementary figures for "Primordial capsid and spooled ssDNA genome structures penetrate ancestral events of eukaryotic viruses"

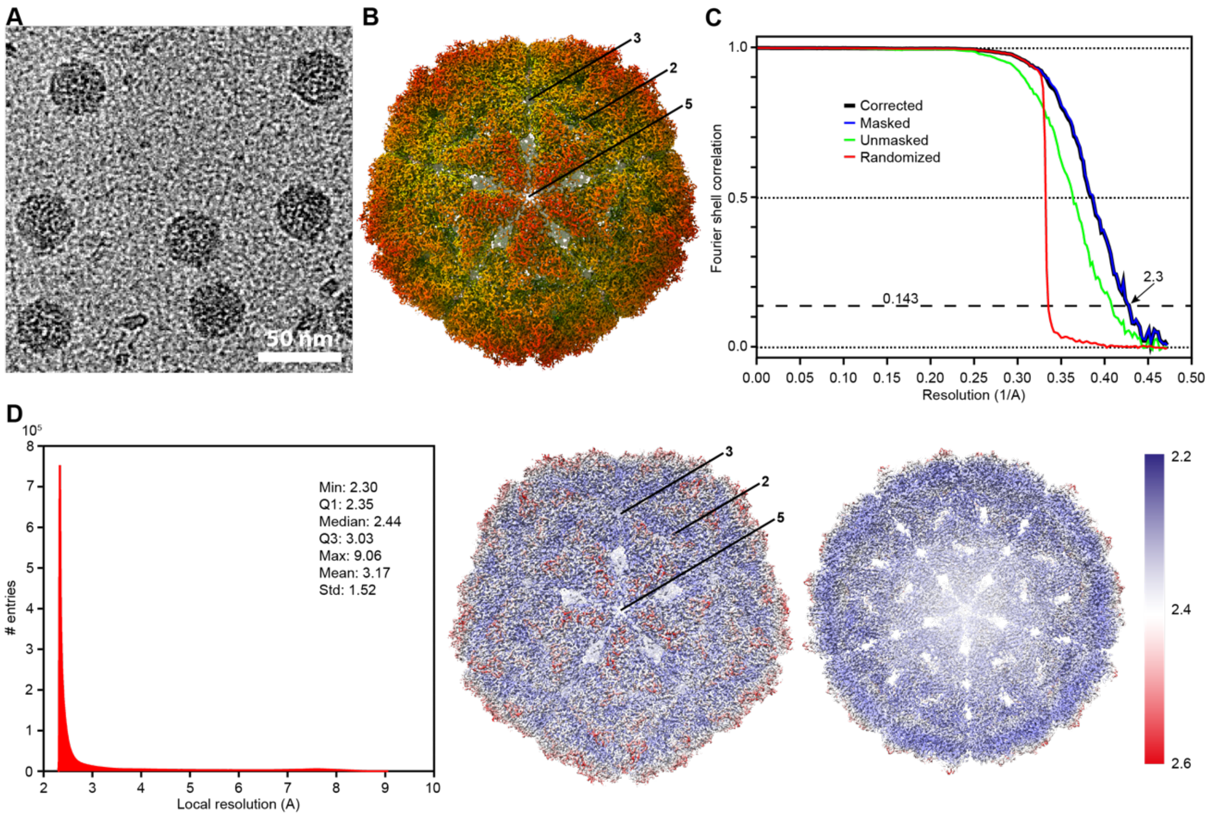

**Supplementary Figure 1. Data collection and reconstruction of the capsid.** (**A**) Cryo-EM raw image of CtenDNAV-II. (**B**) Cryo-EM 3D reconstruction of the CtenDNAV-II capsid. The capsid is viewed down the five-fold axis and radially coloured from blue to red. The icosahedral five-fold, three-fold, and two-fold axes are labelled as 5, 3, and 2, respectively. (**C**) The gold standard FSC resolution curves of masked (blue) and unmasked (green) reconstructions of the CtenDNAV-II capsid. Possible effects of the masking were compensated for by noise randomization (red), to create the final FSC curve (black). The resolution at which the correlation drops below the FSC = 0.143 (gold standard threshold (Henderson et al., 2012; Scheres and Chen, 2012) is 2.3 Å. (**D**) Local resolution of the final reconstruction determined by Relion. The left panel shows a histogram created by Relion showing how many voxels have a given local resolution, and the middle and right panel shows the reconstruction coloured from red to white to blue that corresponds to local resolutions 2.6, 2.4 and 2.2, respectively. The icosahedral five-fold, three-fold, and two-fold axes are labelled as 5, 3, and 2, respectively in the middle panel. The front half of the reconstruction is removed in the right most panel to visualize the inside of the capsid.

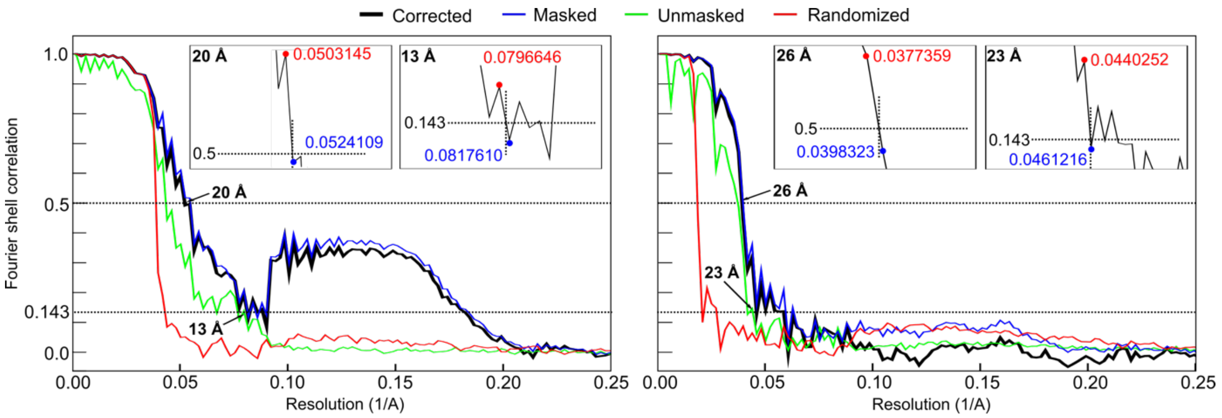

**Supplementary Figure 2. FSC resolution curves of C1 reconstructions.** Left is the FSC curve of the outer layer and right is the FSC curve of the core. The curves and resolution estimates of 13 respectively 23 Å at FSC = 0.143 were generated by Relion. Due to the occasionally noisy curve, the final corrected curve in black at FSC = 0.143 and 0.5 are shown as zoomed in views in the insets, where the circles indicate the two closest x-coordinates.

**Table 1. Data collection and processing**

| | **Capsid EMD-12554** | **Genome – outer layer EMD-12555** | **Genome – core EMD-12556** |
|---|---|---|---|
| Magnification | 140,000 | | |
| Voltage (kV) | 300 | | |
| Defocus range (μm) | -1.00 to -3.00 | | |
| Microscope | Titan Krios | | |
| Camera | K2 | | |
| Total electron dose ($e^-/Å^2$) | 37 | | |
| Pixel size (Å) | 1.06 | | |
| Final particle number | 33,507 | 21,559 | |
| Symmetry imposed | I4 | C1 | |
| Map resolution (Å) (FSC=0.143) | 2.3 | 13 | 23 |
| Map resolution range (Å) | 2.3-9.1 | N/A | |
| Map sharpening B-factor ($Å^2$) | -70.5027 | | |

**Table 2. Structure refinement and validation of capsid.**

| | |
|---|---|
| **Model** | |
| PDB | 7NS0 |
| Composition | |
| Chains | 3 |
| Atoms | 13933 |
| Protein residues | 906 |
| Bonds (RMSD) | |
| Length (Å) | 0.005 |
| Angle (°) | 0.724 |
| MolProbity score | 1.79 |
| Clash score | 5.60 |
| Ramachandran plot (%) | |
| Outliers | 0.00 |
| Allowed | 4.01 |
| Favoured | 95.99 |
| Rotamer outliers (%) | 1.90 |
| Cβ outliers (%) | 0.00 |
| CaBLAM outliers (%) | 1.46 |
| B-factors (Å$^2$) (min/max/mean) | 4.67/33.32/12.03 |
| **Data** | |
| d99 masked (full/half1/half2) | 2.2/3.3/3.3 |
| d99 unmasked (full/half1/half2) | 2.1/3.1/3.1 |
| FSC (model) = 0 (masked/unmasked) | 2.04/2.10 |
| FSC (model) = 0.143 (masked/unmasked) | 2.12/2.25 |
| FSC (model) = 0.5 (masked/unmasked) | 2.32/3.89 |
| **Model vs. Data** | |
| CC (mask) | 0.84 |
| CC (box) | 0.41 |
| CC (peaks) | 0.28 |
| CC (volume) | 0.75 |
| EMRinger score | 6.16 |

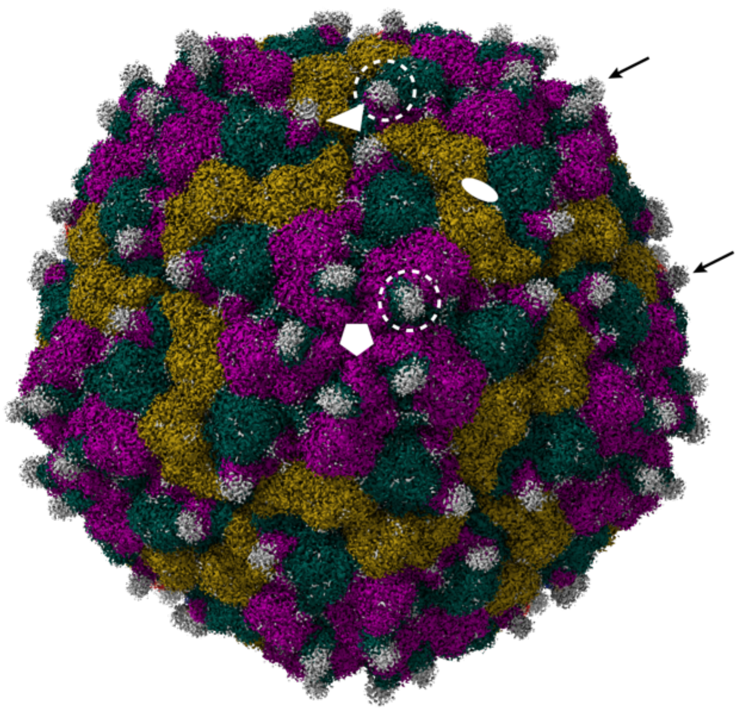

**Supplementary Figure 3**. **Unmodelled density on the capsid surface.** Cryo-EM 3D reconstruction of the CtenDNAV-II capsid at low contour level. The map (originally grey) was coloured according to the model using the command 'color zone' in Chimera X with a colour distance of 6 Å and the same colour code as Fig. 1 (subunit A, purple; subunit B, green; and subunit C, yellow). Unmodelled density from the C-terminal ends of subunit A and B, that remained grey after zone colouring, is visualised around each 3- and 5-fold axis, for which examples are indicated with arrows and dashed circles. The positions of the five-, three-, and two-fold axes are shown by a white pentamer, triangle, and ellipse, respectively.

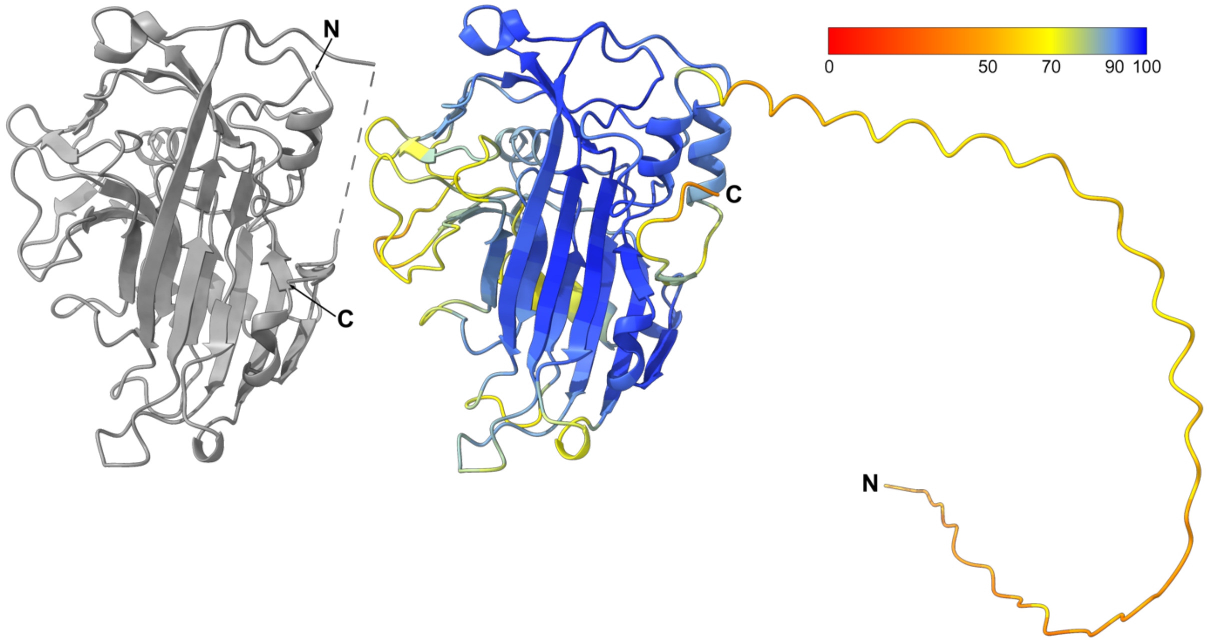

**Supplementary Figure 4. Comparison between the experimentally determined model of the C-subunit and the AlphaFold predicted model.** Left: the experimentally determined model of the C-subunit, right: the computationally predicted structure of the CtenDNAV-II capsid protein using the AlphaFold package. The predicted model is coloured red to blue according to the pLDDT score.

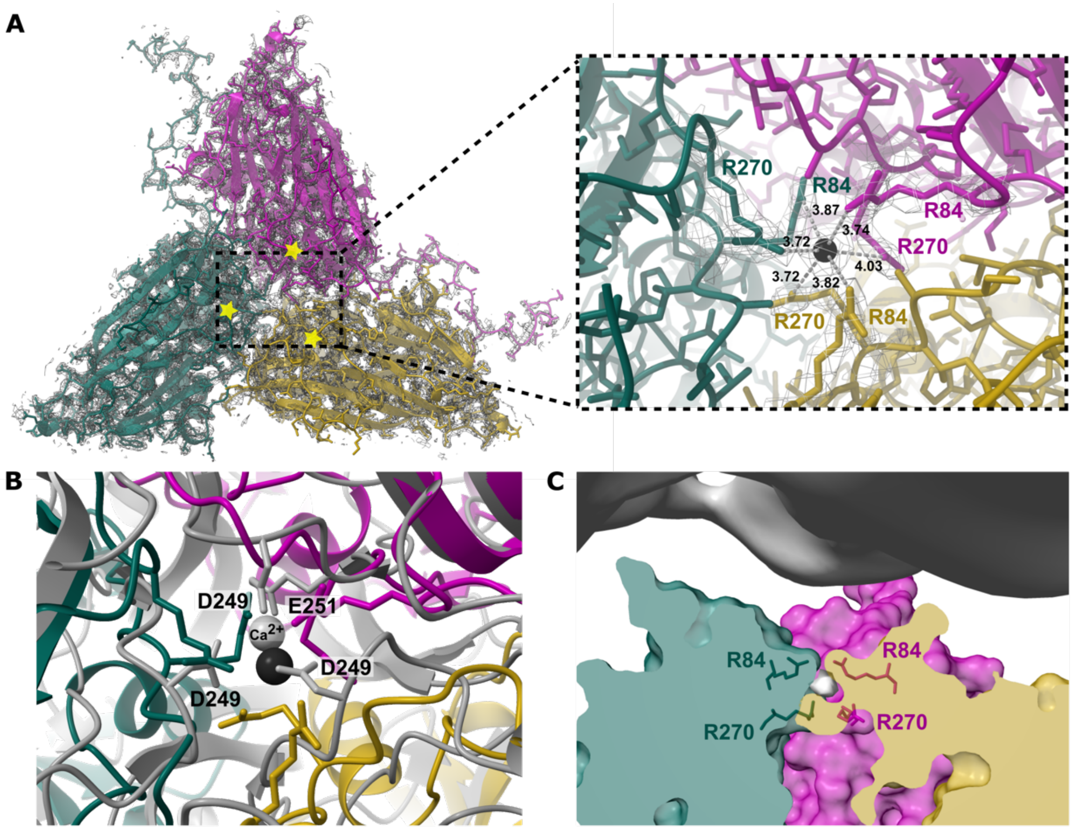

**D**

| | | | | | | | |
|---|---|---|---|---|---|---|---|
| CtenDNAV-II | 77 | QPQIPDG**R**MLESLPR | 91 | 263 | HNNTSAI**R**PKTLQGQ | 277 | N/A |
| YP_004046697.1 | 63 | RPKIPDG**S**TVASLSR | 76 | 249 | LNNNHNI**E**WQDRNQE | 263 | 40% |
| YP_009001776.1 | 69 | QPKIPDG**G**FTSSLSR | 83 | 260 | HPQQTRH**D**PVEIT-K | 273 | 36% |
| AXH78027.1 | 61 | QPKIPDG**E**LTTSLSR | 75 | 261 | HPQSGTH**D**PVELD-L | 274 | 34% |
| YP_004286321.1 | 65 | QPKIPDG**L**LTSSLSR | 79 | 266 | HPRTTTH**D**PTLID-P | 279 | 34% |
| YP_473362.1 | 64 | QPKILDG**A**LTSSLSR | 78 | 260 | HPQSTTH**D**PIILE-Q | 273 | 30% |
| AXH73194.1 | 65 | QPKIPDG**A**FTTSMGR | 79 | 244 | HNISPKD**D**GIINM-Q | 257 | 29% |
| P0DOK2.1 | 72 | QPKIPDG**K**AIMSIGS | 86 | 255 | QAQSEQH**D**FKRIPDR | 269 | 26% |
| QTE03400.1 | 60 | NPKIPDG**K**TYASTGL | 74 | 227 | MPQGNDH**D**FNIIPR- | 240 | 29% |
| QTE03391.1 | 60 | NPKIPDG**K**TYASTGL | 74 | 227 | MPQGNDH**D**FNIIPR- | 240 | 30% |
| QTE03345.1 | 61 | NPKIPDG**K**VYASTGI | 75 | 230 | MPQGNDH**E**FQIIPR- | 243 | 32% |
| YP_009345106.1 | 59 | QPKIPDG**K**VGESVGL | 73 | 235 | RQVKDEV**D**FTQLCSP | 249 | 28% |
| YP_009345096.1 | 58 | QPKIPDG**K**ANESLGF | 72 | 241 | HGQRDYH**D**FIQQRNE | 255 | 27% |
| YP_009109634.1 | 51 | QPKIPDG**K**MTESLGF | 65 | 243 | HGITDEH**D**VKRQQQT | 257 | 24% |
| BBE21063.1 | 50 | RPKIHGD**S**SQLSIGV | 64 | 253 | KHFDGDH**P**CQMIPSR | 267 | 28% |
| QTE03396.1 | 54 | GAKIPDG**K**SYHSTGI | 68 | 228 | MPLGSDH**D**FQRLEQ- | 241 | 27% |
| QKY77697.1 | 57 | QPKIPDG**K**VGTSIGQ | 71 | 234 | HQVKDEV**D**FTQLCTP | 248 | 28% |
| YP_009345085.1 | 59 | QPKIPDG**K**VGTSIGQ | 73 | 236 | HQVKDEV**D**FTQLCTP | 250 | 27% |
| QKY77694.1 | 58 | QPKIPDG**K**VGTSIGQ | 72 | 235 | HQVKDEV**D**FTQLCSP | 249 | 27% |
| AXH76206.1 | 79 | SPKIPDG**K**AELSCGI | 93 | 276 | HPINADH**D**FTETQDS | 290 | 28% |
| QGH73227.1 | 30 | ASKVPDG**K**CGYSIGV | 44 | 214 | QQVNGSR**D**FCTIRQN | 228 | 25% |
| YP_009237569.1 | 20 | TCKIPDG**S**AVLSSSS | 34 | 212 | HREN-DM**D**FVDV--N | 223 | 24% |
| AXH76313.1 | 21 | PMEIKDG**K**GAGSLSL | 35 | 221 | KRTSREH**P**FIGFA-- | 233 | 25% |
| AXH77452.1 | 18 | QPKLLDG**R**VSRTAGI | 32 | 180 | NSTNSEH**E**FVPSEM- | 193 | 23% |

**Supplementary Figure 5. Unassigned density in the subunit interface.** (**A**) The model of a single icosahedral protomer viewed from the inside of the capsid and corresponding cryo-EM map is visualized to the left. To the right, is a close-up view of the subunit interface. A marker (black) was placed in the centre of the unmodelled density using Chimera X to measure the distances to the surrounding arginine residues. The yellow stars mark the N-terminal ends (F64) from each chain. (**B**) A clipped side view of an icosahedral protomer with unmodelled density in grey surrounded by arginine residues. The map from the outer genome layer is shown in the top. (**C**) Same view as in (A), but aligned with Flock House virus (*Nodaviridae*) (PDB 4FTB). Putative amino acids interacting with $Ca^{2+}$ (grey ball) are labelled and shown as sticks. (**D**) Selected regions from a multiple sequence alignment performed on sequences resulting from a BLAST search with default search parameters. The sequence identities to the CtenDNAV-II query sequence are listed to the right. Residues aligned with R84 and R270 are highlighted in bold.

| PDB | Name | Family | Genome | z-score | rmsd | Residues |
|---|---|---|---|---|---|---|
| 2qqp | Providence virus | *Carmotetraviridae* | ssRNA | 15.1 | 2.9 | 209 |
| 3s6p | Helicoverpa Armigera Stunt virus | *Alphatetraviridae* | ssRNA | 14.9 | 2.9 | 208 |
| 4ftb | Flock House virus | *Nodaviridae* | ssRNA | 14.9 | 3.1 | 201 |
| 2bbv | Black Beetle virus | *Nodaviridae* | ssRNA | 14.7 | 2.9 | 199 |
| 1ohf | Nudaurelia capensis omega virus | *Alphatetraviridae* | ssRNA | 14.5 | 3.0 | 207 |
| 1f8v | Pariacoto virus | *Nodaviridae* | ssRNA | 14.5 | 3.2 | 204 |
| 1nov | Nodamura virus | *Nodaviridae* | ssRNA | 14.4 | 3.1 | 202 |
| 3ide | Infectious pancreatic necrosis virus | *Birnaviridae* | dsRNA | 12.1 | 3.0 | 192 |
| 2df7 | Infectous brusal disease virus | *Birnaviridae* | dsRNA | 12.1 | 3.3 | 198 |
| 2izw | Ryegrass mottle virus | *Solemoviridae* | ssRNA | 8.8 | 3.3 | 162 |
| 5zju | Beak and feather disease virus | *Circoviridae* | ssDNA | 8.4 | 4.1 | 155 |

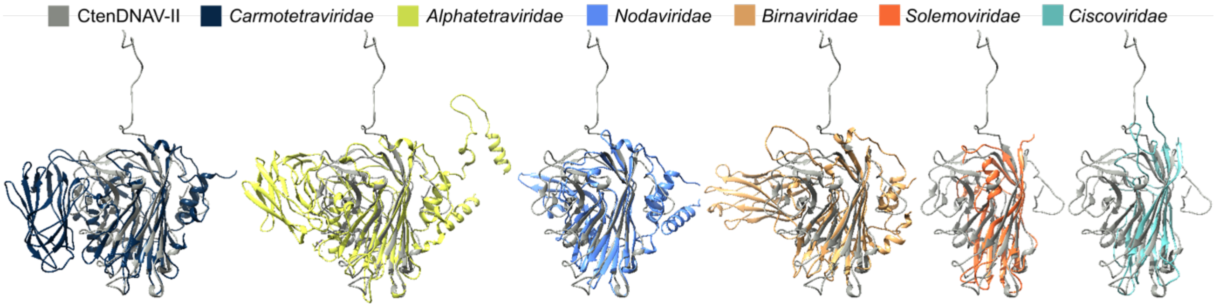

**Supplementary Figure 6. Structural comparison using the DALI server.** A list based on the result from the DALI search showing the 11 most similar viruses, i.e. with highest z-score. For each comparison the z-score, rmsd and the number of residues that were used for the alignment are listed. One representative from each virus family (indicated with *) were used for the structural imposition with the CtenDNAV-II capsid protein (grey).

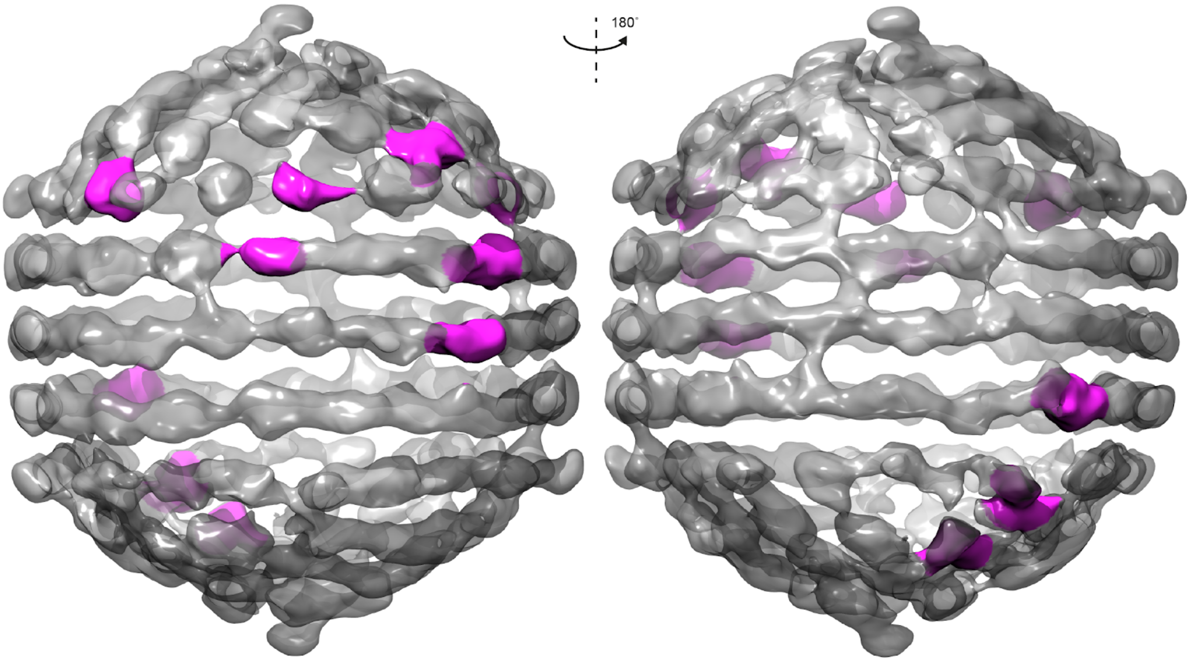

**Supplementary Figure 7. Connections between the outer genome layer and the core.** The outer genome layer where the sites that connect to the core are highlighted in magenta. The connections between the outer genome layer and the core seem to be confined to two specific areas on approximately opposite sides. One area is shown in the left panel and the second is shown to the right, which is rotated 180°.
